# The role of the pre-commissural fornix in episodic autobiographical memory and simulation

**DOI:** 10.1101/706150

**Authors:** Angharad N. Williams, Samuel Ridgeway, Mark Postans, Kim S. Graham, Andrew D. Lawrence, Carl J. Hodgetts

## Abstract

Neuropsychological and functional magnetic resonance imaging (MRI) evidence suggests that the ability to vividly remember our personal past, and imagine future scenarios, involves two closely connected regions: the hippocampus and ventromedial prefrontal cortex (vmPFC). Despite evidence of a direct anatomical connection from hippocampus to vmPFC, it is unknown whether hippocampal-vmPFC structural connectivity supports both past and future-oriented episodic thinking. To address this, we applied diffusion-weighted magnetic resonance imaging (dMRI) and a novel deterministic tractography protocol to reconstruct distinct subdivisions of the fornix previously detected in axonal tracer studies, namely pre-commissural (connecting the hippocampus to vmPFC) and post-commissural (linking the hippocampus and medial diencephalon) fornix, in a group of healthy young adult humans who undertook an adapted past-future autobiographical interview. As predicted, we found that inter-individual differences in pre-commissural - but not post-commissural - fornix microstructure (fractional anisotropy) was significantly correlated with the episodic richness of both past *and* future autobiographical narratives. Notably, these results remained significant when controlling for both non-episodic narrative content and grey matter volumes of the hippocampus and vmPFC. This study provides novel evidence that reconstructing events from one’s personal past, and constructing possible future events, involves a distinct, structurally-instantiated hippocampal-vmPFC pathway.

**Significance Statement:** A novel anatomically-guided protocol that allows the pre-commissural and post-commissural fornix fibers to be separately reconstructed *in vivo* (Christiansen et al., 2016) was applied to reconstruct the pre-commissural subdivision of the white matter fornix tract (anatomically linking the hippocampal formation to the vmPFC) and investigate its contribution to episodic memory and future simulation. We demonstrated that the amount of episodic details contained in past and future narratives, collected via an adapted autobiographical interview, was positively correlated with pre-, but not post-, commissural fornix microstructure. These findings highlight how inter-individual variation in the pre-commissural subdivision of the fornix underpins the construction of self-reflective, contextual events – for both the past and future.

## Introduction

A key adaptive feature of human cognition is the ability to re-experience our personal histories and imagine the future in vivid detail. According to the constructive episodic simulation hypothesis, the processes and neural machinery that allow us to remember past experiences also allow us to imagine future experiences (Schacter et al., 2012; Addis, 2018). Consistent with this view, remembering past and imagining future events activate a common set of brain regions, including the hippocampus and ventromedial prefrontal cortex (vmPFC) (Addis et al., 2007; Benoit and Schacter, 2015). Furthermore, the ability to retrieve episodically rich autobiographical memories and construct coherent future simulations is diminished following lesions to both the hippocampus and vmPFC (Race et al., 2011; McCormick et al., 2018) (but see Squire et al., 2010). Such findings have led to the suggestion that the hippocampus and vmPFC are critical nodes within a default (Andrews-Hanna et al., 2010; Raichle, 2015) or ‘core’ network that interact to support autobiographical memory and imagination (Schacter et al., 2012; Schacter et al., 2017) (For related proposals see also Murray et al., 2017; Robin and Moscovitch, 2017; McCormick et al., 2018).

Converging evidence has shifted focus towards this neural network-level approach to support the way we reconstruct our personal past and construct possible future experiences (Schacter et al., 2012; Bellana et al., 2017; Schacter et al., 2017). For instance, studies using functional magnetic resonance imaging (fMRI) have found increased functional connectivity between the hippocampus and vmPFC during both the retrieval of autobiographical memories (McCormick et al., 2015) and the construction of episodic future events (Campbell et al., 2017), and resting state functional connectivity between these regions has been shown to predict the episodic quality of individual’s memories (Yang et al., 2013).

The communication of information across networked areas depends on the organization and integrity of the white matter connections between them (Jbabdi and Behrens, 2013). Invasive tract-tracing techniques have revealed direct efferent anatomical connections from the hippocampus to the vmPFC. In rats, the entire longitudinal extent of the subiculum/CA1 is connected - via the pre-commissural fornix - with the vmPFC, with connectivity increasing progressively in strength from dorsal to ventral hippocampus (Jay and Witter, 1991; Cenquizca and Swanson, 2007). Similarly in primates, the pre-commissural fornix provides the exclusive route for subiculum/CA1 projections to medial and orbitofrontal PFC (Barbas and Blatt, 1995; Aggleton et al., 2015), with relatively more projections arising from the anterior hippocampus. In humans, diffusion-weighted magnetic resonance imaging (dMRI), which can non-invasively delineate the path of major fiber pathways and evaluate their microstructure through indices such as fractional anisotropy (FA) (Jbabdi and Behrens, 2013), has provided initial evidence for hippocampus-PFC connections via the fornix (Croxson et al., 2005). Building on this work, Christiansen et al. (2016) recently developed an anatomically guided dMRI protocol for the selective *in vivo* reconstruction of pre-commissural fornix fibers in humans, allowing investigation of the functions supported by human hippocampus-PFC structural connectivity for the first time.

By application of this novel anatomically-informed tractography protocol, we investigated the role of the pre-commissural fornix in autobiographical past and future thinking using an individual differences design (Palombo et al., 2018a). Participants were asked to recall past experiences and generate future events using word-cues according to a modified Galton-Crovitz cue-word paradigm (Crovitz and Schiffman, 1974). White matter microstructure was assessed in these individuals using high angular resolution diffusion-weighted imaging (HARDI) and constrained spherical deconvolution tractography (Dell’Acqua and Tournier, 2019). Given the directed hippocampus-PFC functional connections identified above in relation to (re)constructing events in episodic memory and episodic simulation (McCormick et al., 2015; Campbell et al., 2017), we hypothesized that individual differences in the episodic richness of past and future thinking would be related to the microstructure of the hippocampus-PFC connections underpinned by the pre-commissural fornix. As a comparison tract, we used the post-commissural fornix, which connects hippocampus to mammillary bodies and anterior thalamic nuclei (Aggleton, 2012; Christiansen et al., 2016; Mathiasen et al., 2019).

## Materials and methods

### Participants

Participants were 27 healthy Cardiff University undergraduates (aged 18–22 years, mean age = 19, 25 females). Portions of this data have been published previously (Hodgetts et al., 2017a). Here we address a novel and distinct question with unpublished data from a future thinking task and a novel anatomically-informed tractography protocol for reconstructing distinct fornix sub-divisions. Participants completed an adapted autobiographical cue-word paradigm (Crovitz and Schiffman, 1974; Addis et al., 2008) in a separate session from diffusion-weighted magnetic resonance imaging (dMRI). All participants gave written informed consent before participating. Cardiff University School of Psychology Research Ethics Committee reviewed and approved this research.

## Experimental Design

### Past-future Autobiographical Interview (AI) task procedure

Participants completed an adapted autobiographical cue-word paradigm (Crovitz and Schiffman, 1974; Addis et al., 2008) that probed both past and future events. In each of the two conditions (past, future), ten cue-words (e.g. “holiday”, “birthday”) were provided to each participant, in response to which they were asked to recall or imagine a personal event and to generate as much detail as possible within 1-minute (Cole et al., 2012). Each event was required to be spatiotemporally specific, occurring over a timescale of minutes or hours, but no longer than a day. Future events were required to be plausible given the participant’s current plans and not previously experienced by the participant. Three alternate word lists were used; these were matched for semantic category (i.e., participants either heard the cue-word ‘holiday’, ‘journey’ or ‘vacation’). Prior to commencing, participants were instructed:

*“In this test I am going to give you a series of words and ask you to recall an episode from your past, or think of an episode that you might be involved in in the future, related to each of these words. The episode needs to be as specific and detailed as possible. I would like you to give me as much information as you can.”*

In cases where the participant either lacked specificity or detail in their description, the experimenter would provide a non-specific prompt for further information (e.g., *“Is there anything else you can tell me about this event?”).* All trials for one temporal direction (past or future) were completed before beginning the trials for the other condition. Order of presentation of temporal direction (past or future) was counterbalanced, as were the word lists (across the past and future conditions). Participants were tested individually, and responses were recorded using a portable recording device (Zoom H1 Digital Field Recorder) for subsequent transcription and scoring.

### Scoring

The standardized AI scoring procedure (Levine et al., 2002) was used. Events (past and future) generated were segmented into distinct chunks of information in order to allow analysis of the levels of episodic or semantic information provided within each. These chunks were typically characterized by grammatical clauses that referenced a unique occurrence, observation or thought (Levine et al., 2002). Two broad categories were used to categorize details: ‘internal’ details (which described the central event) and ‘external’ details (decontextualized information, including semantic details and information concerning extended events that are not specific in time and place, and repetitions). In the case that a participant described more than one event, the event that was described in the most detail was coded as ‘internal’ and the other as ‘external’. The central event was required to refer to a specific time and place, thus it can be considered ‘episodic’ and will be referred to as such henceforth. Episodic details included not only time and place details, but also any other episodic information (sensory details, thoughts and emotions) that were part of the central event (Levine et al., 2002). **Figure 1** contains examples of external and episodic details from past and future narratives. Total score was computed by summing over the 10 event narratives.

**Figure 1.**
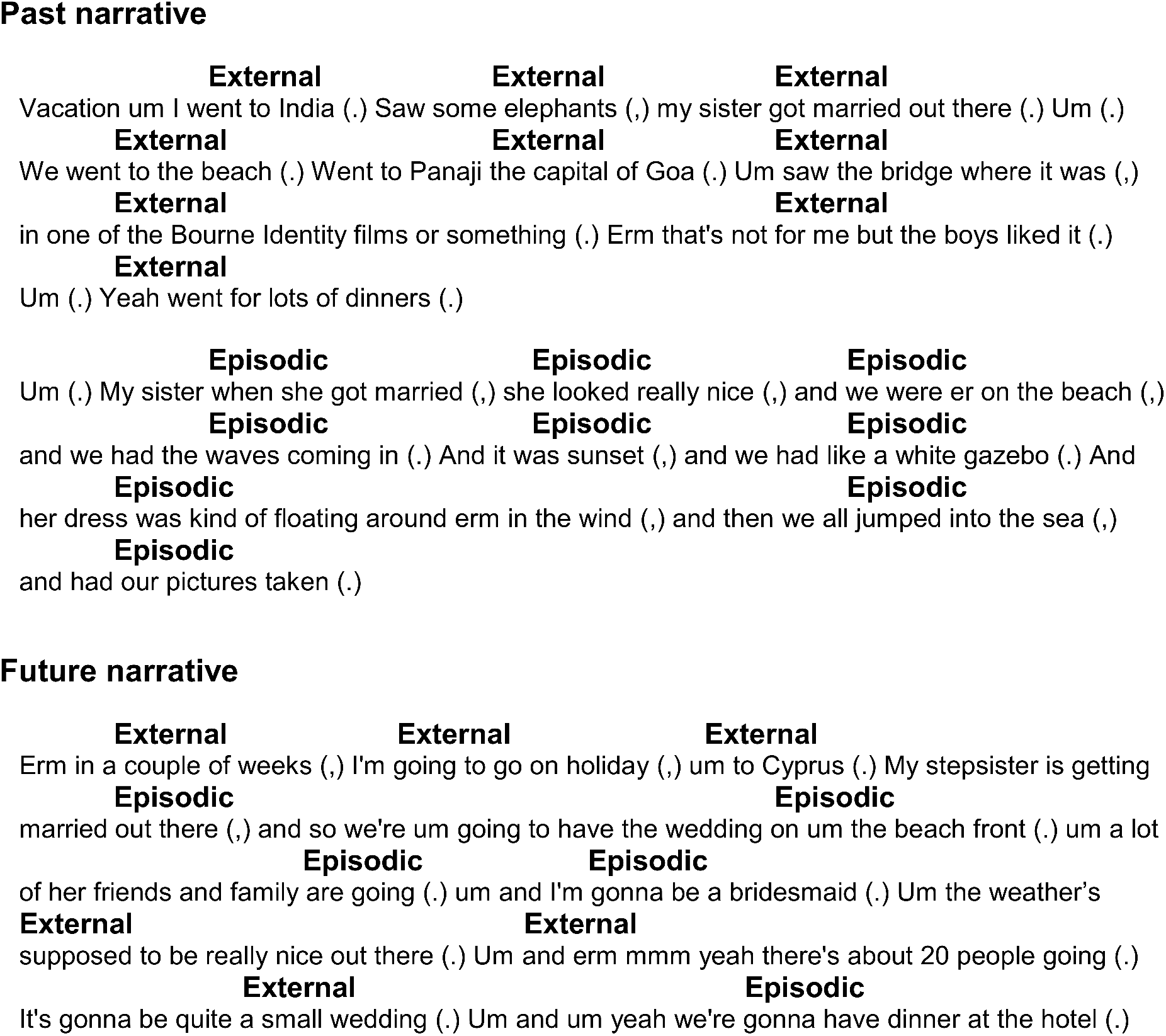
Examples of external and episodic details from past and future narratives.

Consensus scoring was established based on the near perfect inter-rater reliability from two raters who scored the past events (intra-class correlation analysis, two-way random model: episodic r = 0.99; external r = 1.0). The values from one primary coder, who completed both the past and future scoring, were used in the analysis. All raters were blind to dMRI results.

For each event the numbers of episodic and external details were tallied, and the totals were then summed across the 10 events in each condition (past, future) to create episodic and external AI scores for each condition for each participant.

### MRI data acquisition

Imaging data were acquired using a General Electric Healthcare (GE) 3-T HDx MRI system with an 8-channel receive-only head coil, at Cardiff University’s Brain Research Imaging Centre (CUBRIC). A standard T1-weighted 3D FSPGR sequence (178 axial slices, 1mm isotropic resolution, TR/TE = 7.8/3.0s, FOV = 256 × 256 × 176mm, 256 × 256 × 176 data matrix, 20° flip angle) provided high-resolution anatomical images.

A diffusion weighted single-shot spin-echo Echo-Planar Imaging (EPI) pulse sequence was used to acquire whole-brain High Angular Resolution Diffusion Image (HARDI) data (60 contiguous slices acquired along an oblique-axial plane with 2.4mm thickness and no gap, TE = 87ms; voxel dimensions = 2.4 × 2.4 × 2.4mm^3^; FOV = 23 × 23 cm^2^; 96 × 96 acquisition matrix). The acquisition was cardiac gated, with 30 isotropic directions at b = 1200 s/mm^2^. In addition, three non-diffusion weighted images were acquired with b = 0 s/mm^2^.

### MRI preprocessing

#### Diffusion MRI

dMRI data were preprocessed using ExploreDTI version 4.8.3 (Leemans and Jones, 2009). Distortions resulting from eddy currents and participant head motion were corrected. A particular issue for white matter pathways located near the ventricles (e.g., the fornix), is free water contamination from cerebrospinal fluid. This has been shown to significantly affect tract delineation (Concha et al., 2005). Thus, to correct for voxel-wise partial volume artifacts arising from free water contamination, the two-compartment ‘Free Water Elimination’ (FWE) procedure (Pasternak et al., 2009) was applied – this improves Diffusion Tensor Imaging (DTI)-based tract reconstruction and tissue specificity (Pasternak et al., 2014). Following FWE, corrected diffusion tensor indices were computed. Fractional anisotropy (FA) – a DTI-based index proposed to reflect axonal organization (Pierpaoli et al., 1996), reflects the extent to which diffusion within biological tissue is anisotropic (constrained along a single axis) (Beaulieu, 2002). FA values can range from 0 (fully isotropic) to 1 (fully anisotropic). The resulting free water corrected FA maps were inputs for the tractography analysis.

#### Tractography

Deterministic tractography was performed from all voxels based on constrained spherical deconvolution (CSD) (Jeurissen et al., 2011; Dell’Acqua and Tournier, 2019). CSD allows for the representation of bending/crossing/kissing fibers in individual voxels, as multiple peaks in the fiber orientation density function (fODF) can be extracted within each voxel (Dell’Acqua and Tournier, 2019). The step size was 1mm, and the fODF amplitude threshold was 0.1. An angle threshold of 30° was used to prevent the reconstruction of anatomically implausible fibers.

To generate 3D fiber reconstructions of each tract segment, waypoint region-of-interest (ROI) gates were drawn manually onto whole-brain free water corrected FA maps. The waypoint ROIs defined the tracts based on a ‘SEED’ point and Boolean logical operations: ‘NOT’ and ‘AND’. The ‘NOT’ and ‘AND’ gates corresponded to whether tracts passing through were omitted from analyses or retained, respectively. These gates were combined to reconstruct the tracts, based on anatomical plausibility. Initially, a multiple ROI approach was applied to reconstruct the fornix (see Metzler-Baddeley et al., 2011; Hodgetts et al., 2017a).

#### Fornix reconstruction

A ‘SEED’ point ROI was placed on the coronal plane, encompassing the body of the fornix. An ‘AND’ ROI was placed on the axial plane, capturing the crus fornici in both hemispheres at the lower part of the splenium of the corpus callosum. ‘NOT’ ROIs were placed intersecting the corpus callosum on the axial plane, and anterior to the fornix pillars and posterior to the crus fornici on the coronal plane. Further ‘NOT’ way-gates were placed after the initial reconstruction and ensuing visual inspection, to remove anatomically implausible fibers. Subsequently, the anterior body of the fornix was split into the pre- and post- commissural column segments (**Figure 2**).

**Figure 2.**
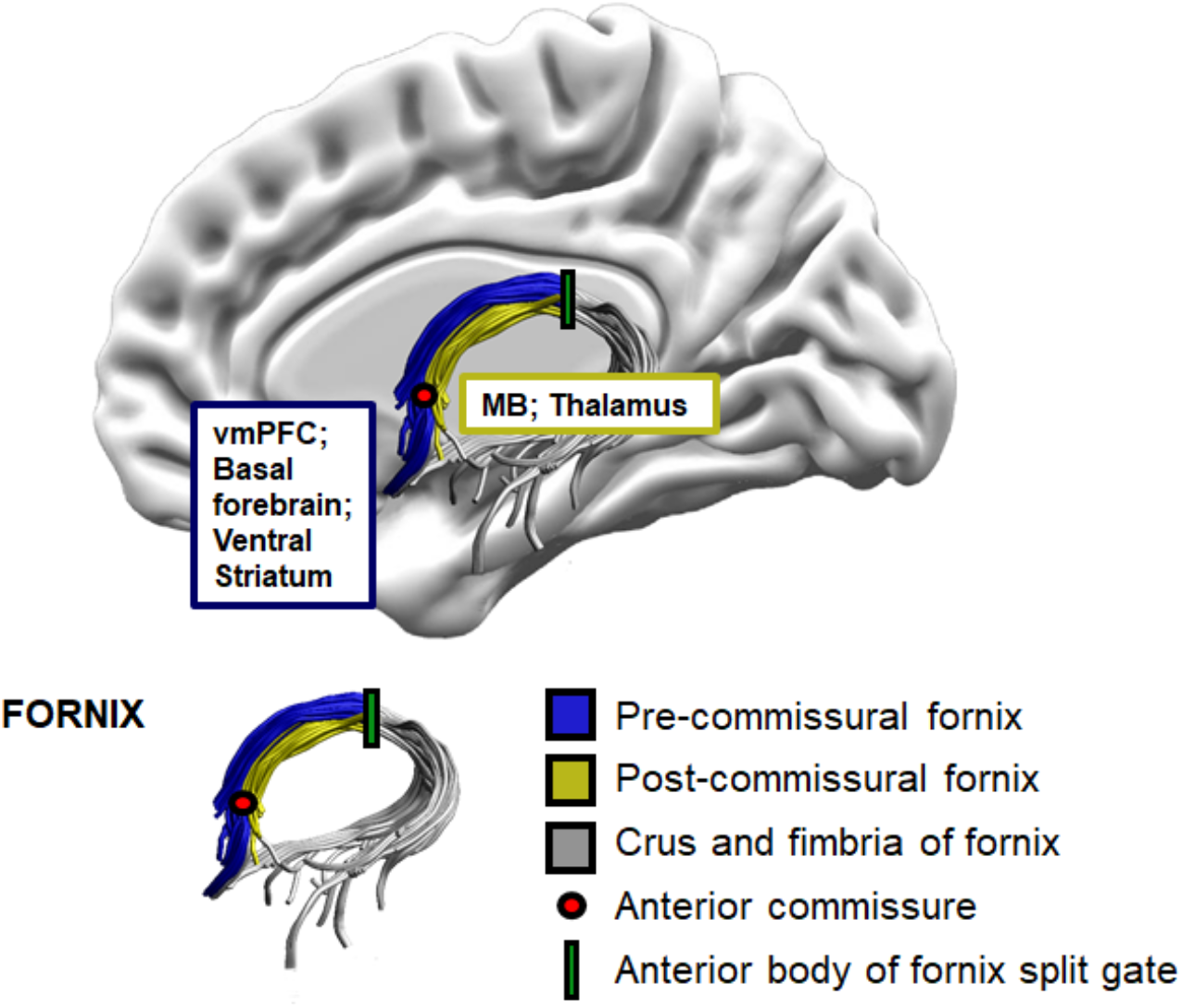
Schematic illustration of the anatomical landmarks for fornix tract sub-division, and the connecting areas of interest. vmPFC = Ventromedial Prefrontal Cortex; MB = Mammillary Bodies.

Waypoint ROIs for the pre-post split (**Figure 3**) were based on the protocol described in Christiansen et al. (2016), and example tract reconstructions are depicted in **Figure 4**. After tract reconstruction for each participant, mean FA values were calculated by averaging the values at each 1mm step along each segment.

**Figure 3.**
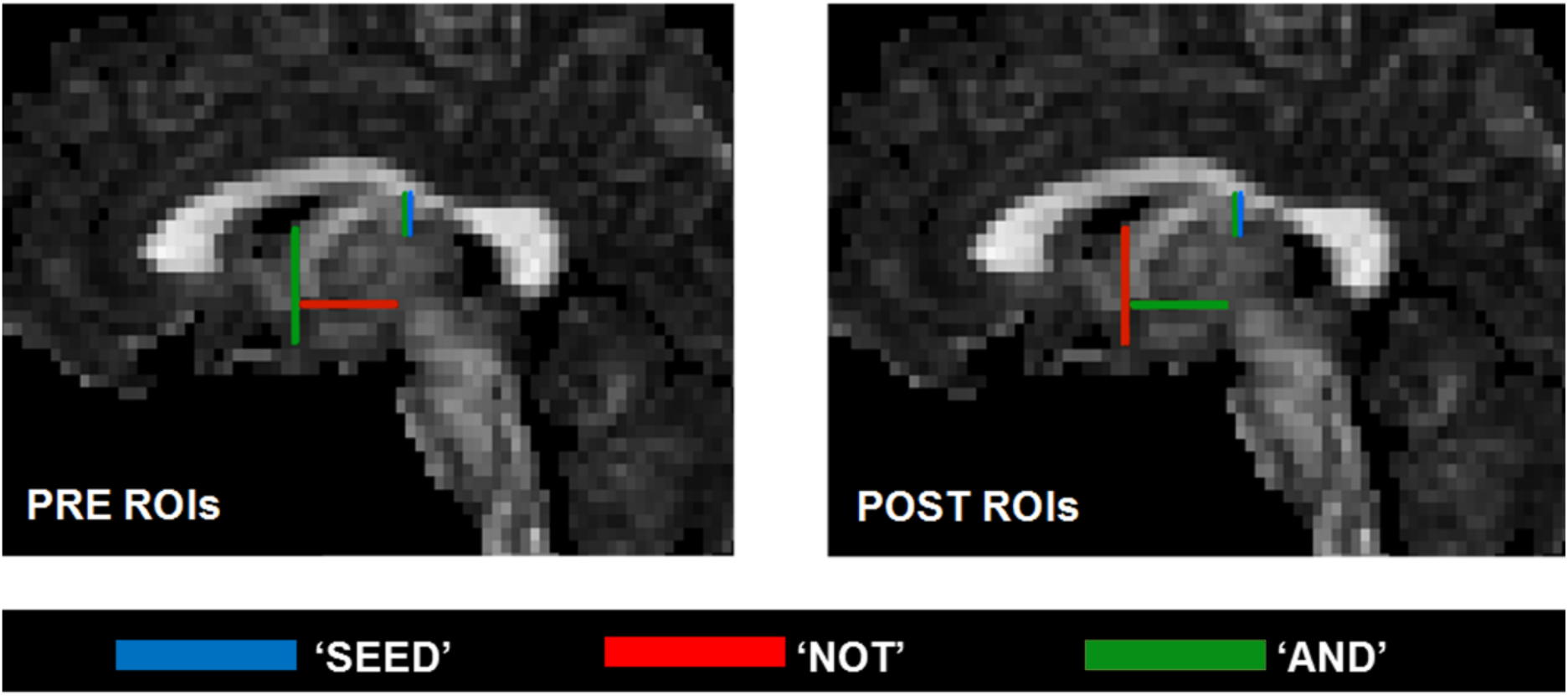
Waypoint region-of-interest (ROI) gates used for reconstructing the pre- and post- commissural fornix tract segments (Blue = SEED, Red = NOT, Green = AND).

**Figure 4.**
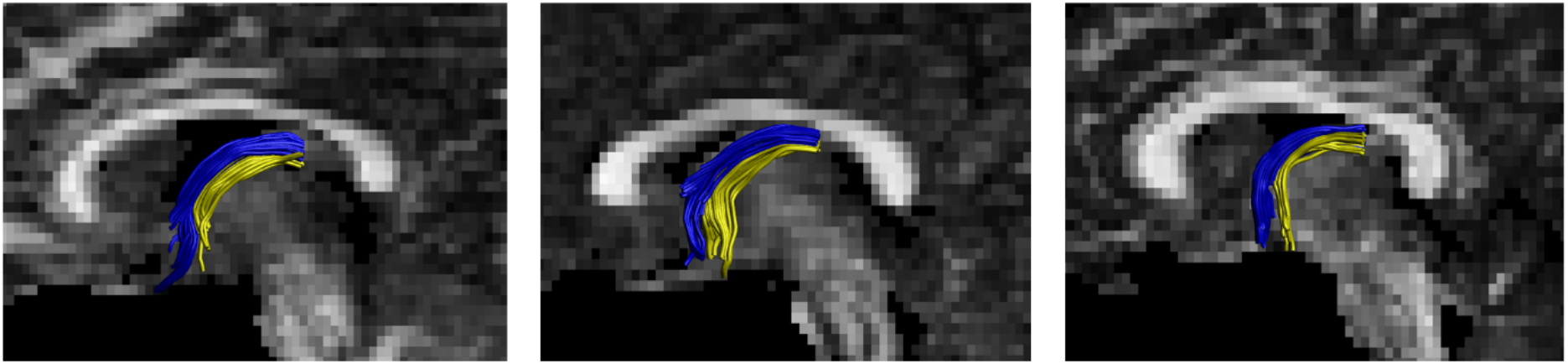
Example reconstructions for the pre- and post- commissural fornix segments (Blue = Pre, Yellow = Post).

#### Pre- and post- commissural fornix reconstruction

The fornix was split, isolating the anterior-body, by an ‘AND’ gate positioned at the point of the downward bend to the crus and fimbria of the fornix. In line with Christiansen et al. (2016), fibers of the crus and fimbria of the fornix were excluded from the anterior-body and hence pre- and post- commissural fornix reconstructions. Partial volume effects due to the intermingling of the two fiber populations beyond the crus were, therefore, minimized (Saunders and Aggleton, 2007). In addition, this procedure avoided ‘jumping’ where tract voxels that pass close to, or across, neighboring tract voxels ‘jump’ onto them (Jones and Cercignani, 2010). This split was conducted using the tract segmentation tool “splitter” within ExploreDTIv4.8.3.

The anterior-body of the fornix was then divided into the pre- and post-commissural segments. This delineation took advantage of the manner in which the fibers separate at the anterior columns of the fornix. At this level, the segments contain approximately the same number of fibers (Powell et al., 1957). The pre-commissural fornix was delineated by positioning an additional ‘AND’ gate on the coronal plane at the anterior-commissure, as well as an additional ‘NOT’ gate meeting this ‘AND’ gate on the axial plane. For the post-commissural fornix reconstruction, the additional ‘NOT’ and ‘AND’ gates placed for reconstruction of the pre-commissural fornix were swopped (see **Figure 3**). Thus, for the pre-commissural fornix, tracts were included only if they extended anterior to the anterior commissure, and for the post-commissural fornix only tracts running posterior to the anterior commissure were retained (see **Figure 4**) (Christiansen et al., 2016).

#### Grey matter volumetrics

T1-weighted images were corrected for spatial intensity variations using FMRIB’s Automated Segmentation Tool (FAST; Zhang et al., 2001). Bilateral grey matter volumes (expressed as a proportion of estimated total intracranial volume) of the hippocampus were subsequently obtained using FMRIB’s Integrated Registration & Segmentation Tool (FIRST; Patenaude et al., 2011). Volumes for the vmPFC ROI were derived using FreeSurfer (surfer.nmr.mgh.harvard.edu: Destrieux et al., 2010), via summing volumes of the medial orbitofrontal cortex (mOFC) and rostral anterior cingulate cortex (rACC) parcels. One participant was removed from the grey matter analyses due to poor overall data quality on the T1 FSPGR.

### Statistical Analysis

As higher values of FA are considered indicative of increased myelination and improved organization, cohesion, and compactness of white matter fiber tracts (Beaulieu, 2002), we predicted a positive association between pre-commissural FA and the episodic richness of past and future constructions. Thus, directional Pearson’s correlations were conducted between individual’s total scores of episodic and external details produced for the ten past and future narratives; and individual’s episodic past and future scores and their FA values for the pre- and post-commissural fornix (Lakens, 2016). Vovk-Sellke Maximum (VS-MPR) *p* ‒ratios were computed: based on the *p* -value, the maximum possible odds in favor of H_1_ over H_0_ equals 1/(−*e p* log(*p*)) for *p* ≤ .37, where log is the natural logarithm and *e* is its constant base (Benjamin and Berger, 2019). I.e. the VS-MPR represents the largest odds in favor of the alternative hypothesis relative to the null hypothesis that is consistent with the observed data (Benjamin and Berger, 2019). Complementary non-parametric Spearman’s rho rank tests were also conducted for the key correlations. These are less sensitive to potential outliers and differences in range (Croux and Dehon, 2010). In addition, partial correlations were conducted for the key episodic-fornix microstructure correlations, to control for the contribution of the number of external details given and regional grey matter volume. All analyses were conducted in JASP (2018, version 0.9.1.0).

## Results

### Correlations between tract microstructure and past-future AI scores

#### Number of details produced (episodic and external) for the past and future narratives

Consistent with previous studies (e.g. Addis et al., 2008; Addis et al., 2009; Race et al., 2011), the total number of episodic details (summed across the 10 cue words) an individual recalled for the past (mean = 121.3, median = 114, SD = 40.8, range = 64 - 247) correlated strongly with the number of episodic details imagined for the future (mean = 59.3, median = 54, SD = 23.4, range = 27 - 105) (**Figure 5A**. Pearson’s r = 0.69, p < 0.001, VS-MPR = 1027.33). Additionally, in line with previous studies, there were significantly more episodic details given for the past in comparison to the future (*t*(26) = 10.75, p < 0.001, d_z_ = 2.07, paired t-test). The number of external details an individual recalled for the past (mean = 73.8, median = 71, SD = 39, range = 20 – 182) also correlated significantly with the number of external details imagined for the future (mean = 86.5, median = 75, SD = 40.8, range = 23 - 198) (**Figure 5B**. Pearson’s r = 0.73, p < 0.001, VS-MPR = 3254.64). There were also significantly more external details given for the future in comparison to the past (*t*(26) = 2.23, p = 0.035, d_z_ = 0.43, paired t-test). The number of episodic details an individual recalled for the past also correlated with the number of external details recalled for the past (Pearson’s r = 0.35, p = 0.035, VS-MPR = 3.15); this was not the case, however, for the future (Pearson’s r = −0.16, p = 0.783, VS-MPR = 1.00).

**Figure 5(A, B).**
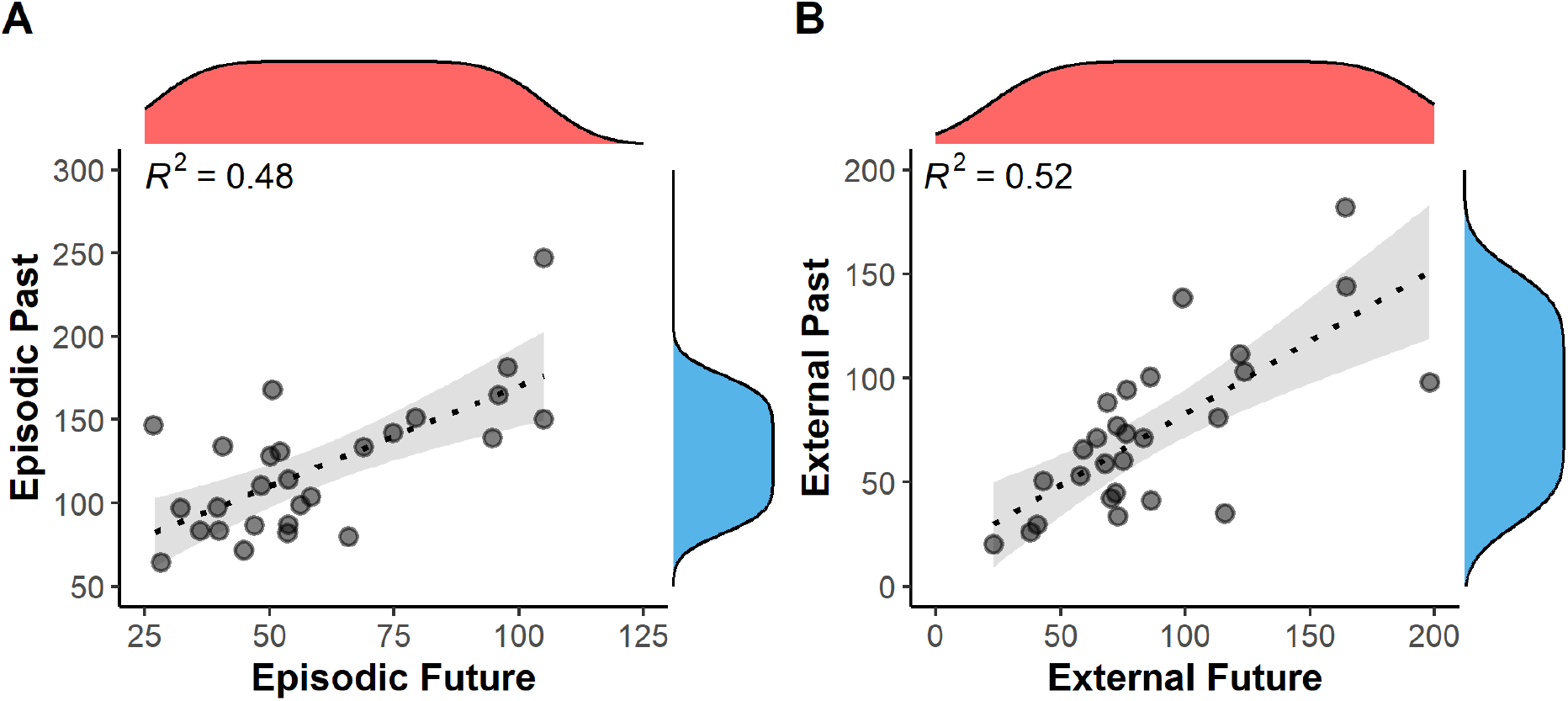
Scatterplots depicting correlations between the number of details produced for the past versus the future AI narratives (A. Episodic, B. External) (*N*=27). Marginal density is displayed on the opposite axis. Grey shading equals the 95% CI.

#### Episodic past details and pre-/post- commissural fornix FA

We found a significant positive correlation between the number of past episodic details and pre-commissural fornix FA (**Figure 6A**. Pearson’s r = 0.49, p = 0.005, VS-MPR = 14.49, Spearman’s rho = 0.464, p = 0.007, VS-MPR = 10.09). There was no significant correlation between post-commissural fornix FA and episodic past details (**Figure 6B**. Pearson’s r = −0.12; p = 0.725, VS-MPR = 1.00, Spearman’s rho = 0.02, p = 0.457, VS-MPR = 1.00). The correlation between episodic past details and pre-commissural fornix FA was significantly greater than between episodic past details and post-commissural fornix FA (Steiger z (27) = 2.29, p = 0.011) (computed using R package ‘cocor’, Diedenhofen and Musch, 2015).

**Figure 6(A-D).**
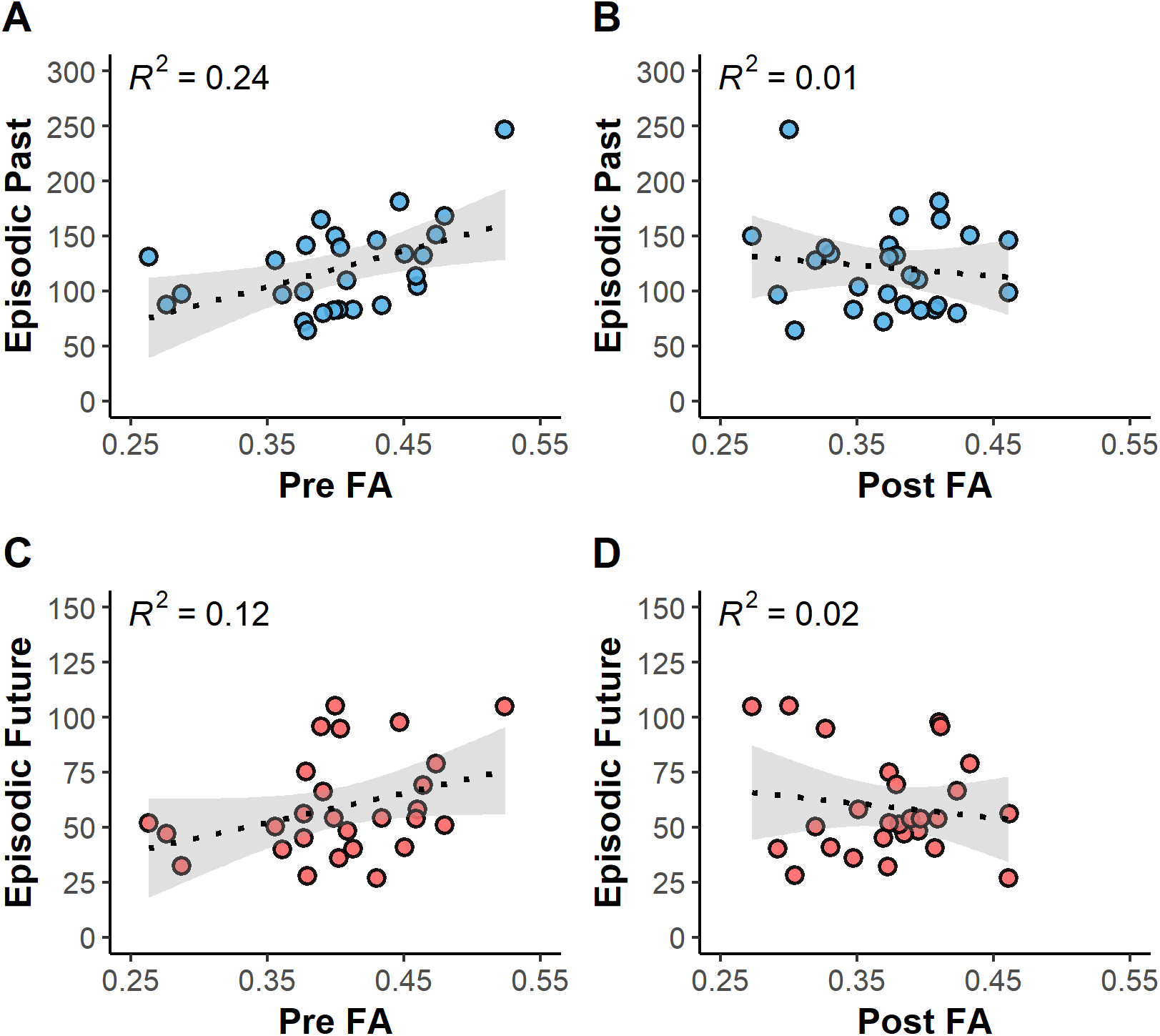
Scatterplots depicting the correlations of episodic past (A, B) and future (C, D) AI details with pre-/post- commissural fornix microstructure (fractional anisotropy, FA). Number of episodic past/future details (summed over 10 cue words) is plotted on the y-axis (*N*=27). Grey shading equals the 95% CI.

The correlation between episodic past details and pre-commissural fornix FA was also significantly greater than between external past details and pre-commissural fornix FA (Steiger z (27) = 1.69, p = 0.046). Additionally, when controlling for the number of external details produced by the individual, the correlation between episodic past details and pre-commissural fornix FA remained significant (Pearson’s r = 0.48, p = 0.007, Spearman’s rho = 0.47, p = 0.007).

#### Episodic future details and pre-/post- commissural fornix FA

The findings for the episodic future simulation details mirrored those for the episodic past retrieval. There was a significant positive correlation between the total number of future episodic details (summed over the 10 cue words) and pre-commissural fornix FA (**Figure 6C**. Pearson’s r = 0.35, p = 0.035, VS-MPR = 3.11, Spearman’s rho = 0.33, p = 0.045, VS-MPR = 2.62), and, correspondingly, there was no significant correlation between episodic future details and post-commissural fornix FA (**Figure 6D**. Pearson’s r = −0.14, p = 0.752, VS-MPR = 1.00, Spearman’s rho = 0.09, p = 0.330, VS-MPR = 1.01). The correlation between episodic future details and pre-commissural fornix FA was also significantly greater than between episodic future details and post-commissural fornix FA (Steiger z (27) = 1.78, p = 0.038). The correlation between episodic future details and pre-commissural fornix FA was not significantly greater than between external future details and pre-commissural fornix FA, however, when controlling for the number of external details given the correlation between episodic future details and pre-commissural fornix FA remained significant (Pearson’s r = 0.38, p = 0.028, Spearman’s rho = 0.33, p = 0.0499).

### Influence of grey matter volume

When hippocampal volume was controlled for, the correlation between past episodic details and pre-commissural fornix FA remained significant (partial correlation: Pearson’s r = 0.49, p = 0.006), and there was no significant association between post-commissural fornix FA and episodic past details (partial correlation: Pearson’s r = −0.09, p = 0.341). Likewise, the correlation between future episodic details and pre-commissural fornix FA remained significant when controlling for hippocampal volume (partial correlation: Pearson’s r = 0.37, p = 0.035), and there was no significant association between post-commissural fornix FA and episodic future details (partial correlation: Pearson’s r = 0.04, p = 0.422).

Similarly, when vmPFC volume was controlled for, the correlation between past episodic details and pre-commissural fornix FA remained significant (partial correlation: Pearson’s r = 0.54, p = 0.003), and there was no significant correlation between post-commissural fornix FA and past episodic details (partial correlation: Pearson’s r = −0.22, p = 0.143). For the future simulations, the correlation between the number of episodic details and pre-commissural fornix FA remained significant when controlling for vmPFC volume (partial correlation: Pearson’s r = 0.36, p = 0.037), and there was no significant correlation between post-commissural fornix FA and episodic future details (partial correlation: Pearson’s r = −0.19, p = 0.178).

## Discussion

We applied a novel anatomically-guided protocol that allows the pre-commissural and post-commissural fornix fibers to be separately reconstructed *in vivo* (Christiansen et al., 2016). To assess both past- and future-oriented thinking, we used an adapted autobiographical cueing paradigm (Crovitz and Schiffman, 1974; Cole et al., 2012) alongside a validated coding scheme that specifically parses episodic from non-episodic detail within individuals’ real-world descriptions (Levine et al., 2002). Using this approach, we found that inter-individual variation in *pre*-commissural, but not *post*-commissural, fornix microstructure was significantly correlated with the amount of episodic detail produced during the construction of both past and future events. Critically, this effect was still seen when controlling for non-episodic content. These findings deepen our understanding of hippocampal-vmPFC interactions in human episodic autobiographical memory and future thinking and provide a ‘structural realization’ of hippocampal-vmPFC functional connectivity (Kosslyn and Van Kleeck, 1990), that is, a direct relationship between the microstructure of the fiber pathway connecting these distributed regions and individual differences in the episodic content of past and future thinking.

Our findings highlight the importance of hippocampus-vmPFC structural connectivity mediated by the pre-commissural fornix (Cenquizca and Swanson, 2007; Aggleton et al., 2015), in episodic construction across past and future events. This builds upon previous fMRI studies that have shown that functional coupling between these distributed regions is increased during both the retrieval of autobiographical memories and the construction of future events (McCormick et al., 2015; Campbell et al., 2017). One recent study, which used structural equation modelling of fMRI data, found increased functional connectivity from anterior hippocampus *to* vmPFC when participants retrieved autobiographical memories in response to cue words (McCormick et al., 2015). Similarly, another investigation applied dynamic causal modeling to fMRI data and found that anterior hippocampus *to* vmPFC effective connectivity increased specifically during the initial construction of episodic future events (Campbell et al., 2017). From this, the authors proposed that ‘the hippocampus initiates event construction in response to retrieval cues, which then drives activation in the vmPFC where episodic details may be further integrated’.

This conceptualization is consistent with previous work in both humans and rodents that demonstrated that hippocampal activity precedes medial PFC activity during memory retrieval (McCormick et al., 2015; Place et al., 2016), and with findings in rodents that hippocampus mediates theta drive to vmPFC (O’Neill et al., 2013). Optogenetic studies in mice (e.g. Ciocchi et al., 2015) have also shown that during memory retrieval ventral hippocampal signals carrying contextual information are sent directly to medial PFC, facilitating coordinated activity between these areas.

The differential contributions of the hippocampus and vmPFC to episodic constructive processes are hotly debated (Robin and Moscovitch, 2017; Schacter et al., 2017; McCormick et al., 2018). According to scene construction theory, the hippocampus, and particularly the subiculum, plays a central role in forming representations of spatially coherent scenes across memory, perception and imagination (Zeidman and Maguire, 2016; Hodgetts et al., 2017b), and these conjunctive scene representations have been proposed to provide a scaffold when constructing both past and future events (Murray et al., 2017; Robin, 2018; Barry and Maguire, 2019). In contrast, the constructive episodic simulation hypothesis contends that the construction of spatiotemporal contexts arises out of a more general relational processing mechanism (Eichenbaum and Cohen, 2014) housed in hippocampus, which is also responsible for the integration of other event details into the event representation (Schacter et al., 2012; Schacter et al., 2017; Addis, 2018). The vmPFC’s contribution to episodic construction, by contrast, has been linked to demands on schematic representations (van Kesteren et al., 2012; Gilboa and Marlatte, 2017; Robin and Moscovitch, 2017), in particular the self-schema (Buckner and Carroll, 2007; D’Argembeau, 2013). For instance, Kurczek et al. (2015) (see also Verfaellie et al., 2019) compared the number of references to ‘the self’ included in autobiographical event narratives from patients with bilateral hippocampal or medial PFC damage as well as healthy controls. Patients with medial PFC damage, despite being able to construct highly detailed episodic events, produced relatively few self-references, and they incorporated themselves in the narratives of their (re)constructions less frequently than the healthy participants. Patients with hippocampal damage showed the opposite pattern: they were impaired in their ability to construct highly detailed episodic events across time periods but not in their incorporation of the self. We have previously suggested (Murray et al., 2017) that hippocampal-vmPFC connectivity serves to (re)create complex conjunctive representations in which one’s self is oriented in a particular time, place, and overall situational context (Murray et al., 2017). These conjunctive representations may subsequently constrain further retrieval and construction by the hippocampus (Graham et al., 2010; Preston and Eichenbaum, 2013; Place et al., 2016; Campbell et al., 2017). Thus, recall/imagination of personally relevant episodes involves a prefrontal system that can work in conjunction with the MTL system to help individuals recombine episodic details to construct a *personally relevant* past/future event (but see scene construction theory - McCormick et al., 2018; Barry and Maguire, 2019; Ciaramelli et al., 2019 - for an alternative account of vmPFC contributions that de-emphasizes self-processes).

Critically, the pre-commissural fornix does not carry reciprocal projections from the vmPFC to the hippocampus (which are indirect via the thalamic nucleus reuniens and entorhinal cortex) (Aggleton et al., 2010; Preston and Eichenbaum, 2013; Murray et al., 2017), but only carries connections to the vmPFC from the hippocampus (subiculum/CA1) (Cenquizca and Swanson, 2007; Aggleton et al., 2015). While several models of episodic memory emphasize the importance of bi-directional interactions between hippocampus and vmPFC (e.g. Preston and Eichenbaum, 2013; Eichenbaum, 2017; Robin and Moscovitch, 2017), with vmPFC playing a regulatory (Preston and Eichenbaum, 2013; Eichenbaum, 2017; Robin and Moscovitch, 2017; Barry and Maguire, 2019) or even initiating (McCormick et al., 2018; Barry et al., 2019) role in episodic construction, our findings reveal that the direct inputs that the hippocampus provides to vmPFC are critical in individual differences for episodic memory and future thinking, and that the pre-commissural fornix is a key link in this broader hippocampal-vmPFC circuit.

While our findings highlight a key role for hippocampal structural connectivity with medial PFC in constructing self-relevant event representations, previous work in humans, primates and rodents has tended to emphasize the importance of connectivity between the hippocampus and medial diencephalon (i.e., mammillary bodies and thalamus) in spatial and contextual memory (Parker and Gaffan, 1997; Aggleton and Brown, 1999), connectivity which is mediated by the *post*- but not the *pre*-commissural fornix (Vann and Nelson, 2015; Christiansen et al., 2016; Mathiasen et al., 2019). While the current findings seemingly challenge this account, one caveat is that our post-commissural fornix tract reconstructions principally involve the connections of the hippocampus with the hypothalamus, including the mammillary bodies, and largely exclude the projections to the anterior thalamic nuclei, as these turn towards posterior regions as the fornix columns descend (Poletti and Creswell, 1977; Christiansen et al., 2016). These thalamic fibers do not form a discrete tract, rather they remain diffuse (Mathiasen et al., 2019). Previous work has demonstrated that thalamic degeneration can impair episodic autobiographical memory and future thinking (Irish et al., 2013). Notably, however, Vann and colleagues (Vann et al., 2011; Vann, 2013) have reported that lesions to the descending post-commissural fornix columns in rats have little impact on spatial memory tests that are sensitive to mammillary body, mammillothalamic tract, anterior thalamic, and hippocampal lesions. The implication of this finding is that the hippocampal-mammillary connection may not be important for all forms of episodic memory, including (as here) those that place demand on constructive and self-referential processing (see also Tedder et al., 2016) (but see Christiansen et al., 2016).

Although FA is highly sensitive to the microstructure of fibers, it lacks biological specificity, and may reflect myelination, axon diameter and packing density, axon permeability and fiber geometry (Jones et al., 2013). Concha et al. (2010), using human DTI-histology comparisons, found that FA of the fornix was strongly positively correlated with axonal membranes (cumulative membrane circumference) and axonal density. Variation in such microstructural properties can influence communication efficiency and synchronicity between distal brain regions (Jbabdi and Behrens, 2013; Pajevic et al., 2014). Future studies using multi-shell diffusion MRI and advanced biophysical modeling to estimate specific microstructural properties including axon density (Assaf et al., 2017) will provide further insight into the specific biological attributes underlying these microstructure-cognition associations. Further, while our sample size was comparable to related investigations (e.g. Postans et al., 2014; Palombo et al., 2018b), and VS-MPRs showed that our findings provide a good level of diagnosticity (Benjamin and Berger, 2019), it will be important to extend our findings to larger lifespan samples.

In summary, we report a novel association between white matter microstructure of the pre-commissural fornix and episodic past and future thinking, thus elucidating a potential anatomical mechanism by which direct hippocampal-to-vmPFC connectivity supports constructive episodic processing. These findings provide important support for the idea of a core-network supporting both the re-construction of past events and the construction of hypothetical events in the future, and that individual differences in structural connectivity may reflect how richly people can reconstruct the past and construct possible futures.

## Acknowledgments

This work was supported by funds from the Medical Research Council (MR/N01233X/1) (KG, AL, AW, MP) and a Wellcome Trust Strategic Award (104943/Z/14/Z) (KG, AL, CH). SR is supported via an ESRC Wales Doctoral Training Partnership PhD studentship. We would like to thank Ofer Pasternak and Greg Parker for providing the free water correction pipeline, John Evans for scanning support, and Naomi Warne for assisting with transcription and autobiographical memory task coding.

